# Cell size control driven by the circadian clock and environment in cyanobacteria

**DOI:** 10.1101/183558

**Authors:** Bruno M. C. Martins, Amy K. Tooke, Philipp Thomas, James C. W. Locke

## Abstract

How cells maintain their size has been extensively studied under constant conditions. In the wild, however, cells rarely experience constant environments. Here, we examine how the 24-hour circadian clock and environmental cycles modulate cell size control and division timings in the cyanobacterium *Synechococcus elongatus* using single-cell time-lapse microscopy. Under constant light, wild type cells follow an apparent sizer-like principle. Closer inspection reveals that the clock generates two subpopulations, with cells born in the subjective day following different division rules from cells born in subjective night. A stochastic model explains how this behaviour emerges from the interaction of cell size control with the clock. We demonstrate that the clock continuously modulates the probability of cell division throughout day and night, rather than solely applying an on-off gate to division as previously proposed. Iterating between modelling and experiments, we go on to show that the combined effects of the environment and the clock on cell division are explained by an effective coupling function. Under naturally graded light-dark cycles, this coupling shifts cell division away from dusk and dawn, when light levels are low and cell growth is reduced. Our analysis allows us to disentangle, and predict the effects of, the complex interactions between the environment, clock, and cell size control.

## Introduction

Organisms control the size of their cells (1–5). In growing cell colonies or tissues they must do this in part by deciding when to divide. The principles of cell growth and division in microorganisms have been studied for many years (6–8). Multiple size control principles have been proposed, including the sizer model, where cells divide at a critical size irrespective of birth size, or the timer model, where cells grow for a set time before dividing (9–15). Recent time-lapse analysis of microbial growth at the single-cell level suggested that many microorganisms follow an ‘adder’ or ‘incremental’ model (16–21), where new-born cells add a constant cell size before dividing again. This principle allows cell size homeostasis at the population level (15, 18).

Although the rules of cell division under constant conditions are being elucidated, cell division in many organisms is controlled by intracellular cues and time-varying environmental signals. For example, cell division and growth are tightly linked to light levels in algae (22–24), while growth is enhanced in the dark in plant hypocotyls (25). The Earth’s cycles of light and dark can thus cause 24-h oscillations in cell division and growth. To anticipate these light-dark cycles many organisms have evolved a circadian clock which drives downstream gene expression with a period of about 24 hours (26). The circadian clock has been shown to be coupled to cell division in many systems, from unicellular organisms (27, 28) to mammals (29–31). It remains unclear how the clock modulates the innate cell growth and the division principles that organisms follow.

The cyanobacterium *Synechococcus elongatus* PCC7942 is an ideal model system to address the question of how cell size homeostasis can be controlled and modulated by the circadian clock and the environment. Cell sizes are easily coupled to the environment as ambient light levels modulate growth (32), which can be monitored in individual cells over time (33–35). An additional advantage is that the key components of the circadian clock in cyanobacteria are well characterised. The core network consists of just three proteins (KaiA, KaiB and KaiC) that generate a 24-hour oscillation in KaiC phosphorylation (36–38). The state of KaiC is then relayed downstream to activate gene expression by global transcription factors such as RpaA (37, 39). Many processes in *S. elongatus* are controlled by its circadian clock (37, 39–41), including the gating of cell division (28, 34, 42). The prevalent idea is that cell division is freely ‘allowed’ at certain times of the day (gate open) and restricted at others (gate closed).

Gating of cell division in *S. elongatus* was first described by Mori et al. under constant light conditions (28). Their results indicated that cell division was blocked in subjective early night, but occurred in the rest of the 24-h day. Single-cell time-lapse studies under constant light conditions have further examined this phenomenon, and suggested a mechanism for it (34, 42). Elevated ATPase activity of KaiC has been proposed to indirectly inhibit FtsZ ring formation through a clock output pathway (34). It remains unclear what effect the coupling of the clock to cell division has on cell size homeostasis for *S. elongatus*, and what are the underlying division rules.

In this work, we examine how the environment and the clock modulate cell size control and the timing of division in *S. elongatus* using single-cell time-lapse microscopy (Fig. 1). Under constant light conditions the clock splits cells into two subpopulations following different size control and division rules. The specific properties of these subpopulations arise from modulation of cell size control by the clock throughout subjective day and night, rather than solely by repressing (gating) cell division in early night. Cells born during subjective night and early subjective day add less length before dividing again, allowing them to divide before the end of the day, whilst cells born during subjective day add more length, avoiding dividing in subjective night. We develop a stochastic model that explains these cellular decisions. To understand the significance of these results, we examine growth and division under realistic graded light-dark (LD) cycles. Combining modelling and experiment, we find that the clock steers division of fast dividing cells away from dusk, and slow dividing cells away from dawn. This prevents cell division from taking place at times when growth arrest could occur due to little or no light (43). Our predictive model reveals the contributions of the circadian clock, environment, and underlying cell size control mechanisms on division throughout the day and night.

**Figure 1:**
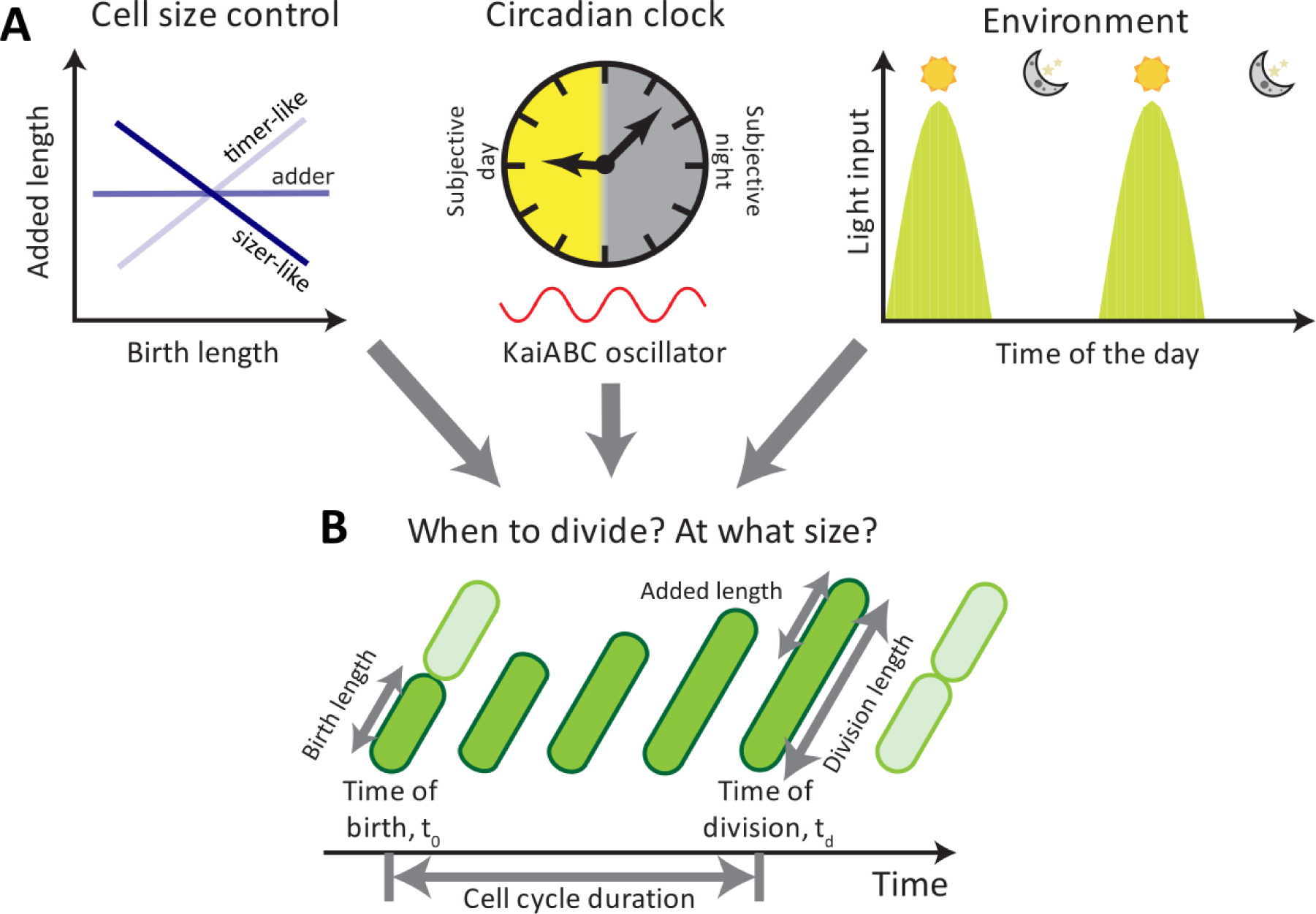
Multiple internal and external factors coordinate cell growth and divisions in cyanobacteria. **(A)**Cell division in cyanobacteria depends on cell-individual factors such as cell size control, the circadian clock, and the environment through light inputs. **(B)** To quantify the impact of these component on a cell’s decision to divide, we quantified timings of birth and divisions and the increase in cell length using time-lapse microscopy.

## Results

### The circadian clock generates two subpopulations following different growth rules under constant light conditions

To examine the role of the clock in cell size control in *S. elongatus*, we first studied growth and division in wild type (WT) and clock-deletion backgrounds under constant light conditions. A clock-deletion strain (Δ*kaiBC*) was obtained by deleting the *kaiBC* locus, thus inactivating the KaiABC oscillator. We carried out time-lapse movies under constant 15 μE m^−2^ s^−1^ cool white light, and segmented and tracked thousands of individual cell lineages over multiple generations. The relation between size at birth and size added between birth and division is often indicative of the model controlling when cells decide to divide (15, 18). If size at birth is linearly related to added size with a slope of 1, then the underlying model is called a 'timer’, in which cells wait a specific time before division. A slope of −1 is indicative of a ‘sizer’, where cells divide after reaching a critical size. More generally negative slopes can be categorised as sizer-like while positive slopes represent timer-like strategies (15, 44–46). Alternatively, added size may not correlate with size at birth (slope of 0). Such cells, which grow by a fixed size, irrespective of their birth size, are described as ‘adders' (16–18) (Fig. 1). *E. coli* and other microbes have been shown to obey this adder rule (15). *S. elongatus* cells are rod-shaped and grow in volume by increasing their pole-to-pole length, and so cell length is an appropriate measure of cell size (SI Sec. 1). Interestingly, in the WT background, *S. elongatus* cells are best fit by a sizer-like model (slope of −0.63), where the larger they are when born, the less length they need to add to reach a target length (Fig. 2A). This effect was less apparent in the clock-deletion background, where cells appeared to have a much weaker dependence on birth length (Fig. 2B) (slope of −0.35).

**Figure 2:**
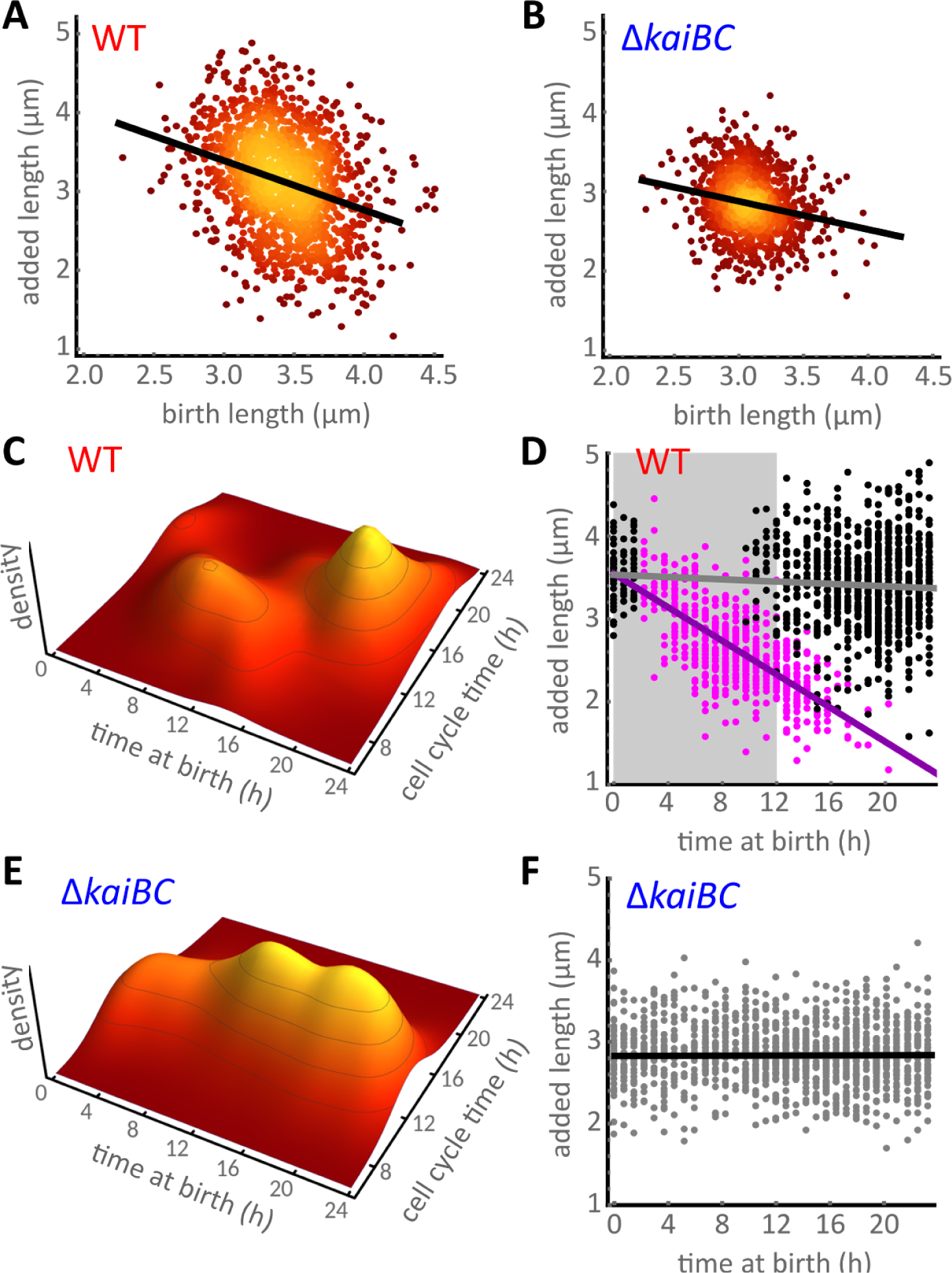
Coordination of cell size control by the circadian clock in *Synechococcus elongatus* generates two subpopulations under constant light. **(A)** Wild type cells (WT) follow an apparent sizer-like principle, where cells add shorter lengths the longer their birth length is (black line, linear regression with slope of −0.63). 1529 individual cells from 3 independent experiments were analysed. Lighter colour indicates a higher density of points. **(B)** Cell size control in a clock-deletion mutant (Δ*kaiBC*) follows an adder-like principle more closely, with a weaker dependence of added length on birth length (black line, slope of −0.35). 1348 individual cells from 3 independent experiments were analysed. Colourmap as in (A). **(C)** The dynamics of WT cells exhibits two distinct subpopulations: cells born late in subjective day have longer cell cycles than cells born earlier. **(D)** Clustering of the two subpopulations reveals an anti-correlation (slope of −0.1 μm h-1, violet line) between time of birth and added length in cells born in late subjective night or early subjective day (magenta dots). These cells also add less length than the subpopulation of cells born during subjective day (black dots). Violet and grey lines show linear regression of the two subpopulations. The shaded grey area represents subjective night. **(E-F)** In populations of the clock-deletion cells, cell cycle times and added length do not depend on the time of day at birth. Black line in (F) is the linear regression line.

How can the circadian clock, which times processes to particular times of the 24-hour day, cause cells to divide at a specific size? To address this question, we first examined how cell division is affected by the time of day. As has been reported previously (28, 34, 42) we observed apparent gating of cell division, with fewer cell divisions in the early subjective night in the WT but not in the clock-deletion background (Supp. Fig. 1). We next asked how this imbalance affects cell cycle times. The distribution of cell cycle times was not clearly bimodal (Supp. Fig. 2), but by clustering cells based on subjective time of birth and cell cycle time (SI Sec. 3 and 4), we found cells lie in two distinct subpopulations (Fig. 2C). WT cells born either in late subjective night or early subjective day have shorter cell cycles (‘fast’ cells) than those born later in the day (‘slow’ cells).

Finally, the timing of cell division also affects added length. On average, cells born in late subjective night or early subjective day add less length (magenta dots in Fig. 2D), as expected from their shorter cell cycle times. Interestingly, within this subpopulation added length decreases with time of birth (violet line, Fig. 2D), allowing these cells to divide before the end of subjective day. This suggests that the clock does not solely enable an off gate at the beginning of night as previously proposed, but it actively promotes cell division to occur before the end of the day. By contrast, in the absence of the clock, no two subpopulations are apparent (Fig. 2E). This is because in the clock-deletion strain cell cycle duration does not depend on the time of birth, and added length is constant throughout the day (Fig. 2F).

### A simple model explains the coordination of cell size by the circadian clock

How does the clock generate the observed complex relationship between added cell length and time of day? To answer this question, we developed a simple model assuming a linear dependence Δ(*L*_0_) = *a L*_0_ + Δ_0_ between added length Δ and birth length *L*_0_. The birth-length independent part of added length Δ_0_ is a stochastic variable. The parameter *a* quantifies the dependence of added length on birth length, and can be estimated by linear regression. Similar models have been used to quantify cell size control of microbes (15, 44, 45).

In *S. elongatus* the circadian clock affects the size control (Fig. 2). We assume that the clock alters the length-independent part of added length Δ_0_ = *L* − (1 + *a*) *L*_0_ by modulating the probability of cell divisions throughout the day. This model assumes that the cellular division rate depends on three factors: (i) increase in cell length 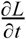, (ii) cell size control *S*(Δ_0_), and (iii) a coupling function *G*(*t*) imposed by the circadian clock, a periodic function of the time of the day *t*, leading to

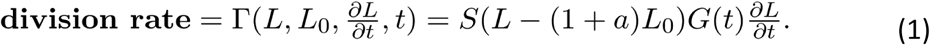

The division rate thus depends on the instantaneous length, length at birth, growth rate and time of the day.

To systematically disentangle the individual components affecting cell division rate, we measured the length-independent part of added length Δ_0_ in clock-deletion cells, which do not gate or modulate cell divisions throughout the day ( *G*(*t*) =1). We found Δ_0_ is well fitted by a Gamma distribution (Fig. 3A, blue line), implying *S*(Δ_0_) is an increasing function of Δ_0_ (Fig. 3A, black line, see SI Sec. 6 for details). Simulations of the resulting stochastic model (Materials and methods) recover the correct size control in clock-deletion cells (Fig. 3B).

**Figure 3:**
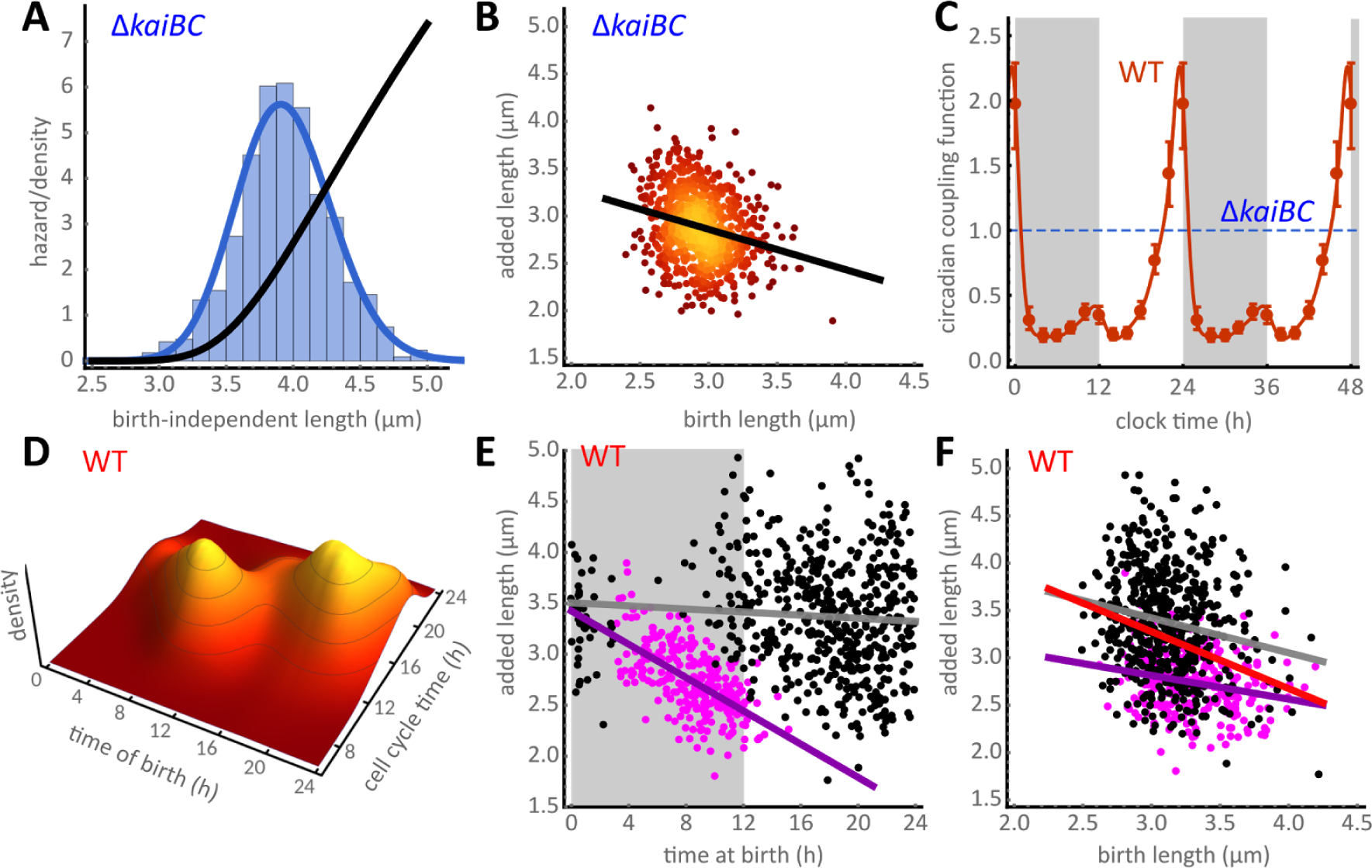
A simple model explains the emergence of the two subpopulations. Cell division rate is coordinated by growth rate, size control and by the circadian clock. **(A)** Distribution of the birth-length independent added length (Δ_0_, blue bars) is fit by a Gamma distribution (blue solid line). The corresponding cell size control hazard *S*(Δ_0_) increases with length (solid black line). **(B)** Stochastic simulations of the model reproduce the size control in the clock-deletion strain (slope of −0.35, compare with Fig. 2B). Lighter colour indicates a higher density of points. **(C)** Bayesian inference from single-cell traces reveals circadian modulation of cell divisions throughout the 24-hour day in the WT. The coupling function *G*(*t*), parameterized by a positive periodic spline (red line, error bars denote 95% credible intervals at knots), decreases the division probability during subjective night (shaded grey background) and early subjective day (white background), when compared to the clock-deletion background (dashed blue line). The coupling coupling then peaks towards the end of subjective day. This function is depicted for two cycles to highlight periodicity. **(D)** Simulations using the inferred coupling function predict the emergence of two subpopulations in close agreement with experimental data (Fig. 2C). **(E)** Clustering of the simulated data (subset shown) also predicts added length to decrease during the day in the fast subpopulation (violet line, slope −0.1 μm h-1) but not in the slow population (grey line). Shaded grey area indicates subjective night. **(F)** The model predicts larger added cell lengths in the slow subpopulation (black dots) compared to clock-deletion cells, and smaller added cell lengths in the fast subpopulation (magenta dots). The respective slopes, which quantify the dependence of added length on birth length in the two subpopulations, are −0.37 (grey line) and −0.25 (violet line). The difference in cell size of the two populations explains the strong dependence of added length on birth length seen in WT cells (red line, slope −0.60).

We then used the model to estimate the circadian coupling function *G*(*t*) directly from individual cell length traces of WT cells via Bayesian inference. The method is based on the likelihood of divisions, which can be obtained analytically from Eq. (1) and is a function of the cell length history (SI Sec. 7). To avoid prior assumptions on the functional form of the coupling, we only constrained it to be a smooth, positive and periodic function of the time of day. Our analysis reveals that the circadian coupling function (Fig. 3C, red line) is at a low basal level throughout subjective night and early subjective day, effectively delaying cell divisions. Then, the coupling function progressively increases peaking towards the end of subjective day, where it facilitates cell divisions compared to the clock-deletion strain (Fig. 3C, dashed blue line).

We also found that elongation (growth) rates oscillate in a circadian manner, which is not apparent in the clock-deletion background. This highlights that the circadian clock not only affects the decision to divide but it also feeds back on growth (43, 47). To account for these effects, we measured the mean trend of these oscillations throughout the day (Supp. Fig. 3, SI Sec. 2) and incorporated the time-dependent growth rate α(*t*) into the WT model. Although the mean trend may average out growth dynamics apparent at the single cell level (47), it greatly simplifies the following analysis.

To predict the WT behaviour, we developed an exact simulation algorithm to carry out detailed stochastic simulations of the model with the inferred coupling function (Materials and methods). In agreement with the experiments (Fig. 2C), the simulations reveal the emergence of differentially timed subpopulations with respect to their birth times (Fig. 3D). We then asked whether the model could also explain the differences in size control observed in the two subpopulations. By clustering the simulation data, we verified that added length decreases throughout the day in the fast subpopulation (magenta dots, Fig. 3E) but not in the slow population (black dots, Fig. 3E), in close agreement with the experiments (Fig. 2D). In contrast, when we forced the coupling function to take the form of a classical on-off gate, whether a piecewise square function (34) or a smoother continuous function (42), we were unable to capture these features of our data (Supp. Fig. 4).

Interestingly, cells in the fast subpopulation modify their effective cell size control to more closely conform with the adder principle (violet line, Fig. 3F), but those in the slow subpopulation follow a weak sizer-like trend similar to clock-deletion cells (grey line, Fig 3F), in excellent agreement with experiments (Supp. Fig. 5). The stronger dependence of added length on birth-length seen in the overall WT population is thus an emergent phenomenon (red line, Fig. 3F, Supp. Fig. 5). It arises from the differentially timed and sized subpopulations generated by the circadian clock.

### Environmental light-dark cycles combine with the clock to generate an effective coupling function

Like all other photoautotrophs, *S. elongatus* did not evolve under constant light. We therefore examined the effects of the circadian clock on growth and division under conditions more relevant to the natural environment of cyanobacteria. We grew WT and clock-deletion cells under graded 12 hour light and 12 hour dark cycles (12:12 LD) that approximate the Earth’s cycles of daylight and dark (Materials and methods, Fig. 4A,B). We programmed the light levels such that the flux of light per unit area over a period of 24 hours is identical in constant light and in 12:12 LD.

**Figure 4:**
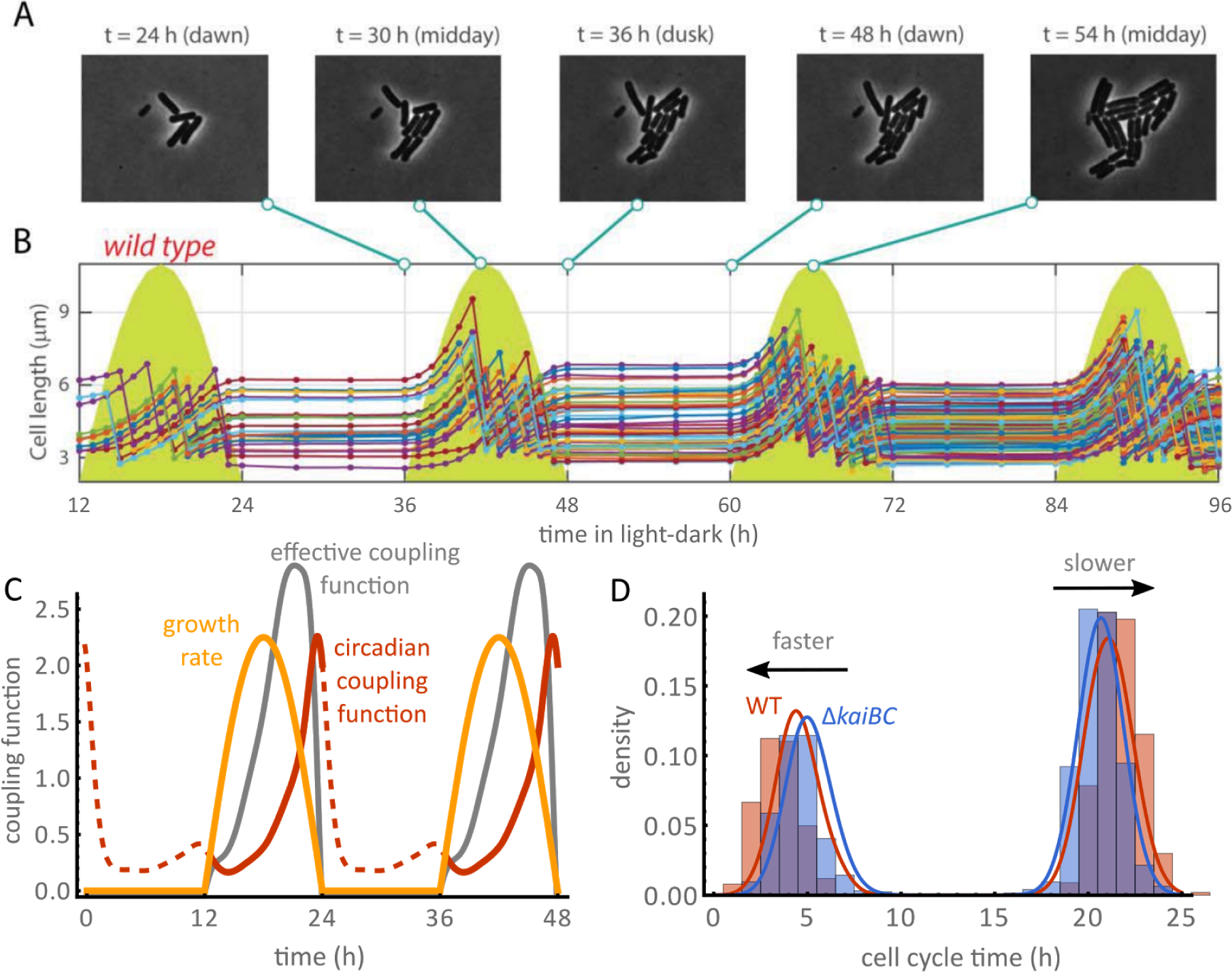
The rate of growth under light-dark (LD) cycles is determined by the light profile for WT and clock-deletion cells, and modifies the coupling function. **(A)** Phase contrast images of a WT micro-colony under graded 12:12 LD cycles reveal periods of growth followed by periods of stagnation between dusk and dawn. **(B)** Length profile of single cells (coloured lines) in a time-lapse movie. Cell growth and divisions are suspended in the dark and slowly rise when the lights turn on. The yellow shades indicate the light profile imposed on cells during the experiment (maximum at midday is approximately 47 μE m^−2^ s^−1^). **(C)** Growth rates of WT (yellow line) and clock-deletion cells closely follow the imposed light profiles (Supp Fig. 6). Growth rate (yellow) and circadian clock (constant light coupling function in red) impose an effective coupling function (grey line, Eq. 2) shifting the probability of cell division towards the end of the day. **(D)** The light conditions split cells into two subpopulations: those that complete a full cycle when the lights are on (short cell cycle times), and those that must wait until the next day to complete a cell cycle (long cell cycle times). Our model shows that the fast WT population (red line) is faster than in the clock-deletion mutant (blue line), but the slow subpopulation is faster in the clock-deletion background. This prediction is confirmed by experimental histograms of cell cycle times for WT (1009 cells from two experiments) and clock-deletion cells (1201 cells from two experiments).

There was no visible growth or cell division in the dark (Fig. 4A,B). As such, the pattern of LD cycles controls the growth rate of both WT and clock-deletion cells forbidding cell divisions in the dark. Elongation rates are also set by the level of ambient light during the day. In graded LD cycles the mean elongation rates of the two strains are nearly identical, and both track the level of ambient light quite accurately (Supp. Fig. 6, Fig. 4C). Restricting growth also constrains the distribution of cell cycle times, separating cells into two subpopulations: cells that divide in the same day they were born (fast cells) and cells that divide only the day after they were born (slow cells) (Fig. 4D). This effect was observed in both WT and clock-deletion cells.

We therefore wondered how the circadian clock interacts with growth cycles imposed by the ambient light levels. By imposing the time-dependent growth rates onto our model of WT and clock-deletion cells, we found that the clock accelerates cell divisions in the fast subpopulation but, interestingly, it delays divisions in the slow subpopulation (blue and red lines, Fig. 4D). In qualitative agreement with this result, we found experimentally that cell cycle times of the two subpopulations were shifted in opposing directions (blue and red bars, Fig. 4D).

These findings are explained through an effective coupling function that results from the interaction of entrained growth rate with the circadian clock. Assuming exponential elongation with rate α(*t*), the effective coupling in our model is given by the daytime-dependent part of the division rate,

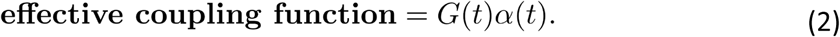

In comparison to clock-deletion cells, effective coupling delays cell divisions at dawn but accelerates divisions close to dusk in the WT (Fig. 4C), highlighting the predictive power of our model. In the following sections, we answer the question of what role the clock plays for size control in light-dark cycles.

### The circadian clock modulates cell size in light-dark cycles

To understand the effect of varying light levels on cell size control, we used the model to predict the type of cell size control in the two subpopulations in LD cycles. Our model predicted that WT cells with short cell cycles, i.e. cells that are born and divide within the same day (magenta dots in Fig. 5A,C, upper panels), add roughly half the length of cells with longer cell cycles, i.e. cells that divide a day after they were born (black dots, Fig. 5A,C, upper panels). Whereas in constant light, slow cells, with longer cell cycle times, grow larger on average, such a supposition is not necessarily true in LD conditions. This is because slow cells typically also live through the lowest light levels, i.e. the least favourable conditions. Indeed, no difference in added length between the two subpopulations is predicted for the clock-deletion mutant (Fig. 5B). The dependence of added length on subpopulation type and the differences between the two strains are well confirmed by the experiments (Fig. 5A,B, lower panels).

**Figure 5:**
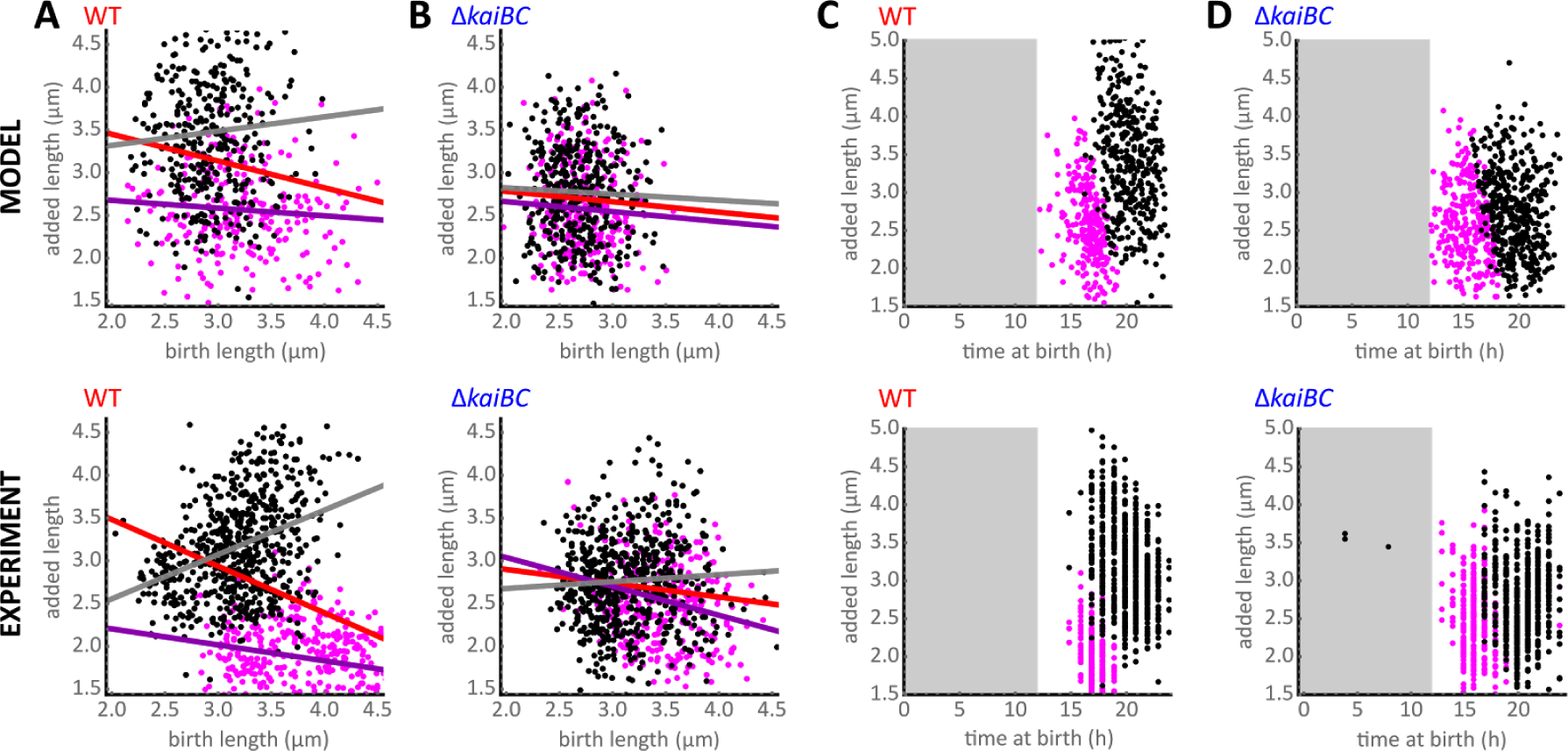
The circadian clock modulates the size control in 12:12 LD cycles. **(A)** The model (upper panel) predicts that the two subpopulations of cells in the WT (short cell cycles in magenta and long cell cycles in black) obey different cell size control strategies. Fast dividing cells obey an adder-like principle (violet line from linear regression, slope of −0.09) while slowly dividing cells show a timer-like size control (grey line, slope of +0.17), which is well confirmed by the experiments (lower panel, violet line slope of -0.19, grey line slope of +0.52). Red line is the overall trend when ignoring the presence of two subpopulations. **(B)** In the clock-deletion population, slow and fast cells obey similar trends in size control compared to the overall population. **(C)** Fast cells (magenta dots) are born earlier in the day on average, and add less length than cells whose cell cycle spans to the next day (black dots). **(D)** In contrast to the WT, in the clock deletion mutant, both subpopulations increment approximately the same length to their birth lengths. Grey shaded areas in (C-D) indicate darkness (lights off).

Furthermore, the model suggests that cell size controls obeys different rules in the two dynamical subpopulations. Fast dividing cells increment their length with a weak dependence on birth size (Fig. 5A, violet line), ie. an adder-like size control. On the other hand, added length of slow cells increases with birth size (grey line), ie. a timer-like size control. This prediction was also confirmed by experiments (Fig. 5A, lower panel). clock-deletion cells, on the other hand, do not display significant differences in cell size and control between the two subpopulations (Fig. 5B,D).

### The effective coupling of divisions to the environment and the clock modulates the frequency of cell divisions in light-dark cycles

We next asked whether the clock affects the time at which cells are born in graded light-dark cycles. In the absence of a clock, *G*(*t*) =1, and so ambient light effectively dictates the division rate (Eq. (2), Fig. 6B, Supp. Fig. 6B). In WT cells, however, our model predicts fewer cell divisions at dawn but more divisions in the middle of the day, which results from the profile of the effective coupling function (Eq. (2), Fig. 6A). This prediction is confirmed by the distributions of time at division (Fig. 6B, SI Sec. 5). WT cells do not divide immediately after dawn, and wait longer than cells in the clock-deletion strain (Fig. 6B). Specifically, we find that 90% of WT cells divide within a window of 4 to 10 hours after dawn as compared to 2 to 10 hours for the clock-deletion mutant, in quantitative agreement with the model.

**Figure 6:**
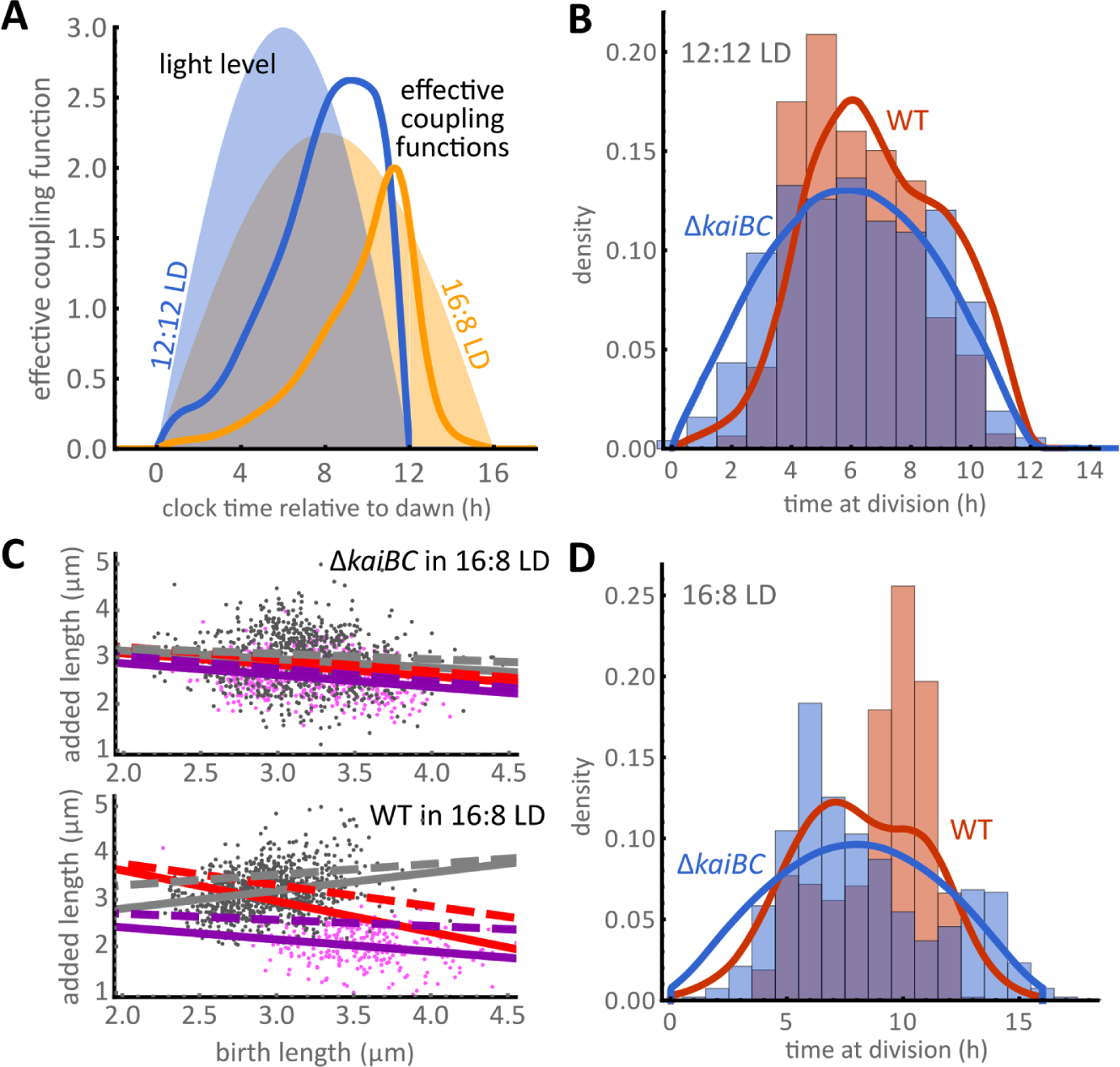
The circadian clock steers cell divisions away from dawn and dusk. **(A)** Imposed light levels (shaded areas) interact with the circadian clock to create an effective coupling function (solid lines). For 12:12 LD cycles the effective coupling function suggests that cell divisions occur away from dawn, while in 16:8 cycles divisions are shifted away from dawn and dusk. **(B)** Division time distributions of WT (red) and clock-deletion cells (blue) obtained from experiments in 12:12 LD (bars) match model predictions (lines). **(C)** In 16:8 LD cycles, WT cells exist in two subpopulations with short (magenta dots from experiments) and long (black dots from experiments) cell cycles that obey different cell size control strategies. Fast dividing cells follow a sizer-like principle (solid violet line), while slowly dividing cells follow a timer-like size control (solid grey line), a dependence also predicted by the model (dashed lines). Red lines are the overall trends when ignoring the presence of two subpopulations. All lines are linear regressions. **(D)** Experimental distribution of division times (bars obtained from 1226 WT and 1372 clock-deletion cells, two experiments each) and model predictions (lines) in 16:8 LD cycles.

To test our understanding of this effect, we interrogated the model under prolonged light durations. In graded 16:8 LD cycles the effective coupling function suggests that the presence of the clock would cause fewer divisions at dawn and dusk, as this effective function peaks closer to midday (Fig. 6A). To test this prediction, we repeated the experiment in a 16:8 LD condition. First, we confirm that the clock affects cell size control similarly to the 12:12 LD condition (Fig. 6C). In WT and clock-deletion backgrounds alike, cells exhibit slow and fast dividing subpopulations. However, owing to the clock, it is only in WT that these two subpopulations exhibit different cell sizes and size control rules (Fig. 6C). The experiments further confirm that WT cell divisions occur in a much narrower range of the day (90% of cells divide between 5 and 12 hours after dawn, Fig. 6D) than for the clock-deletion mutant (4 and 14 hours after dawn). In effect, most cells divide closer to midday in agreement with the theoretical conclusions drawn from the effective coupling function (Fig. 6A).

## Discussion

In this work, we used single-cell data and systematic interrogation of a stochastic model of cell growth to elucidate how the circadian clock and the environment rework underlying rules of cell size control in *S. elongatus* (Fig. 1). We first characterised cell size control in constant conditions and found that the clock generates an apparent sizer-like behaviour (Figure 2A). We showed that this effect is an epiphenomenon caused by the clock generating two subpopulations following different division rules in WT cells (Figure 2C,D, Supp. Fig. 5). These subpopulations differ in cell cycle duration, added size, and times of birth and division relative to a 24 hour day, while no such coordination was present in clock-deletion cells (Fig. 2E,F). These results show that organisms possessing clocks, or coupling their cell cycle to intracellular or extracellular processes that drive them out of steady-state, could have complex size control rules. These can even include more than a single type of cell size control for the same growth condition.

We then formulated a phenomenological model of how the circadian clock coordinates cell size control and cell division rate. The model confirmed that this interaction indeed generates two differently timed and sized subpopulations. Statistical inference using constant light data revealed that the clock modulates cell divisions by progressively increasing the division rate just before subjective dusk, while decreasing it at other times of the 24-h day (Fig. 3C).

Under graded LD cycles, the model predicted that the clock accelerates divisions in the fast subpopulation but delays divisions in the slow subpopulation, a finding that we confirmed experimentally under natural 12:12 LD cycles. By doing so, the clock constricts the time window of cell division. This effect was even more apparent, in both the model and experiments, under a 16:8 LD cycles condition, where divisions are driven away from dawn and dusk. Such a modulation of the timing of cell divisions could provide a fitness advantage by, for instance, avoiding cell division during the energetically unfavourable periods around dawn and dusk. Previous work suggested that the circadian clock’s slowing of growth rate before dusk can aid individual cell survival (48). In future, it will be important to investigate whether the clock’s restriction of cell division towards the middle of the day plays a similar functional role.

By inferring the coupling function between the clock and the cell cycle under constant light conditions, we revealed the qualitative features of clock control of cell size in *S. elongatus*. The clock progressively increases the probability of division throughout the second half of subjective day. The probability of division reaches a well-defined peak, two-fold higher than the clock-deletion reference, just before dusk, before dropping to a basal level after dusk. This adds to our understanding of how the clock controls the cell cycle, revealing that the probability of division is under continuous circadian regulation.

Previous studies have proposed that the clock gates cell division by imposing an off gate in the early hours of subjective dark, and thus causing the scarcity of cell division events observed during that window (28, 34, 42). In our model, this scarcity is generated by the sudden relaxation of the coupling function back to its basal level following the “rush” of divisions before dusk (Fig. 3C), which can be interpreted as an effective gate. However, the peak in the coupling function also generates a progressive decrease in added size towards subjective dusk, as observed experimentally. This effect is not predicted by a classical two-level (on and off) gating function (Supp. Fig. 4). Our finding thus explains the dependence of cell size on the time of birth.

Remarkably, the coupling function, fitted just on constant light data, accurately predicts the effects of the clock on cell size in both 12:12 and 16:8 light conditions. These predictions include the reduction of cell cycle times for cells that divide in the same day they were born, and the increase of cell cycle times for cells that divide in the next day. We elucidate that these complex phenomena can be understood through an effective coupling function accounting for the clock and environment. Our understanding of these non-trivial effects could help to reveal the clock’s function in controlling cell division. In this respect, it will be critical to understand the molecular mechanism behind this coupling function in future work. One possibility is that the mechanism could be a combination of the repressive effects of KaiC ATPase activity on the cell cycle (34) with circadian activation of cell cycle control genes. For example, FtsZ expression is under circadian regulation, peaking near dusk (49). In summary, by incorporating statistics of both cell size and division times, our findings shed light on how the circadian clock governs a cell’s decision to divide at different sizes.

Although our simple coupling function reproduces most of the qualitative features of the cell size control and division rules observed experimentally, in future it will be interesting to examine the few aspects our model could not explain. For example, in clock-deletion cells under 16:8 LD cycles, the division frequency does not seem to follow the light levels imposed throughout the day (blue bars, Fig. 6C). Instead, the distribution of division times is bimodal. This phenomenon could either be caused by an entrainment effect delaying cell divisions at dawn, or be due to a dependency of cell size control ( *S* in Eq. 1) on the light level, which our model does not describe. Further work is needed to elucidate these effects in greater detail. Also, in this work we did not take into account that the clock phase can be modulated by light dark cycles (48).

Examining the relationship between added cell size and birth size has provided valuable insights into how microbes maintain cell size (15–18, 21). However, cells are subject to internal or external cues, which can affect cell size control in non-intuitive ways. In other organisms, including higher eukaryotes, cell division is also subject to a range of internal and external inputs, including the circadian clock. In future, it will be critical to develop predictive models of how these inputs feed into the regulation of cell physiology in these other organisms, similar to what we have done here. As we showed, such models provide unprecedented insights by disentangling the components affecting cellular decision making.

## Materials and methods

### Strains, plasmids and DNA manipulations

*Synechococcus elongatus* WT was obtained from an ATCC cell line (ATCC^®^ 33912^TM^). A clock-deletion strain was generated by insertion of a gentamycin resistance cassette into the ORF of the *kaiBC* operon. The plasmid (a gift from Prof. Erin O’Shea), carrying the interrupted gene with the antibiotic selection marker, was transformed into the WT strain through homologous recombination. Complete allele replacement on all the chromosomal copies was checked through PCR.

**Table 1:**
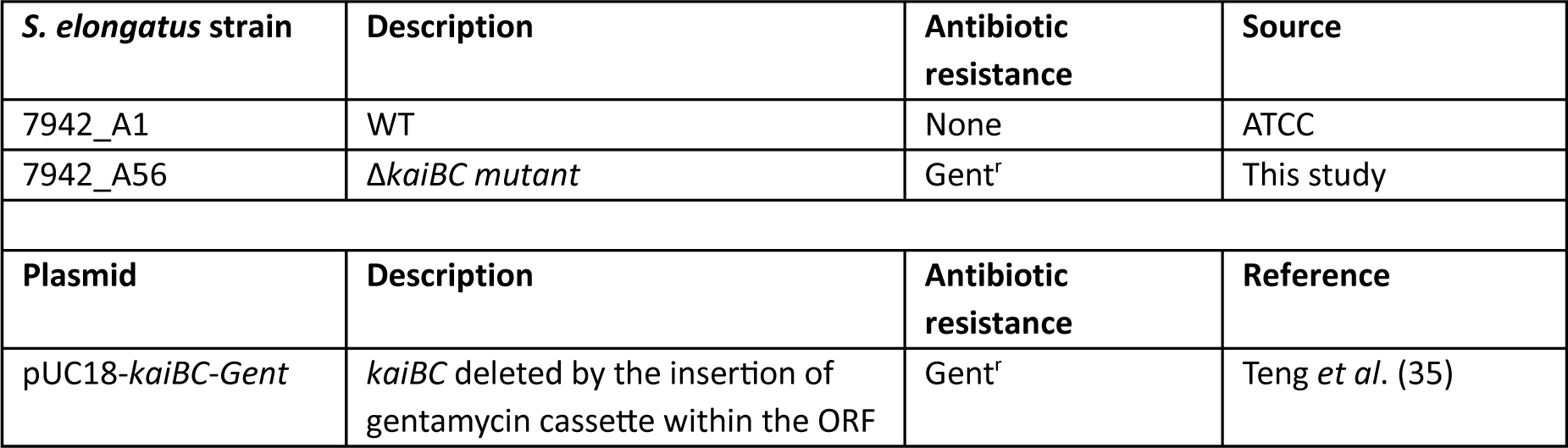
Strains and plasmid used in this study.

### Growth conditions

The strains were grown from frozen stocks in BG-11 media at 30°C under photoautotrophic conditions with constant rotation. The Δ*kaiBC* strain was supplemented with gentamycin at 2 μg ml^-1^. Light conditions were maintained at approximately 20-25 μE m^−2^ s^−1^ by cool fluorescent light sources. Before the start of each movie acquisition the cultures were kept at exponential phase, and entrained by subjecting the cells to a 12 h light: 12 h dark cycle (12:12 LD).

### Microscopy and sample preparation

A Nikon Ti-E inverted microscope equipped with the Nikon Perfect Focus System module was used to acquire time-lapse movies of the cells over several days. 2 μl of entrained cultures in exponential phase were diluted to an OD_750_ of 0.15-0.20 and spread on agarose pads. The agarose pads were left to dry and then placed inside a two chambered coverglass (Labtek Services, UK), which was brought under the microscope. The cells were maintained in a 12:12 LD regime for another cycle, and image acquisition was only started after dawn. Illumination for photoautotrophic growth is provided by a circular cool white light LED array (Cairn Research, UK), attached to the condenser lens of the microscope. Light conditions were pre-programmed to run during acquisition. The setup allows for instantaneous and near-continuous light level updates. In constant light experiments, the light level was set at approximately 15 μE m^−2^ s^−1^. In experiments with light-dark cycles, light was set such that the flux of photons per unit area is the same as in constant light over a 24 hour period. The daily profile of solar insolation in the wild was approximated by the function

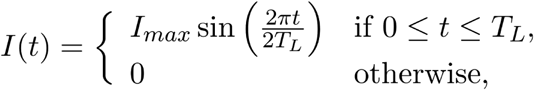

where *T_L_* is the duration of the light period (12 h or 16h), and 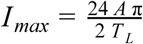. *A* ≈ 15 μE m^−2^ s^−1^ is light level in constant light, and so *I_max_*≍ 47 μE m^−2^ s^−1^ in 12:12 LD. Data acquisition was controlled through the software Metamorph (Molecular Devices, California). At each time point, phase contrast and fluorescent images using the filter set 41027-Calcium Crimson (Chroma Technology, Vermont) and a CoolSNAP HQ2 camera (Photometrics, Arizona) were acquired. The fluorescent image was acquired with low exposure time and low excitation intensity in order not to disturb cell growth, and was used to improve image segmentation. Further, we did not examine cells carrying fluorescent reporters in order to avoid any potential effects of phototoxicity on growth rates, and thus on cell size. In constant light, images were acquired every 45 minutes. In light-dark conditions, images were acquired every hour during the day. The reduction in the frequency of acquisition was implemented to avoid phototoxicity when the light levels are very low and growth is slow. The frequency of acquisition was therefore further reduced in the dark. Finally, the light levels were updated after each stage position is visited and acquired. In between time points, the light levels were updated every 2 minutes. All experiments used a 100X objective. This protocol was adapted from Young *et al.* (50).

### Exact stochastic simulation algorithm coupling the circadian clock to cell size control

We provide an exact simulation algorithm to simulate a lineage of an exponentially growing and dividing cell from *t*_0_ to *T* with division rate Γ (Eq. (1)). The simulation uses a thinning method (51, 52) with a time-horizon Δ*t* over which the division rate Γ is bounded by a constant *B*. For this purpose, we assume that (i) the function *S* is monotonically increasing, and (ii) the coupling function and exponential growth rate are bounded by *G_max_* = *sup G*(*t*) and α*_max_* = *sup* α(*t*), respectively.

#### Algorithm

**Initialize** *t ← t_0_. L(t) ← L_0_*.

**While** *t ≤ T*:

1. Simulate *L(t)* from *t* to *t + Δt* using 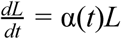. (cell growth)

2. Draw an exponentially distributed random variable τ with rate *B = S(L(t + Δt) − (1 + a)L_0_) G_max_ α_max_L(t + Δt)*.

3. **If** *τ > Δt*:

Update *t ← t + Δt*.

4. **Else**:

Update *t ← t + τ*.

Evaluate Γ(t) and draw a uniform random variable *u*.

**If** *Γ(t) ≥ B u*:

Update *L(t) ← L(t)/2* and *L_0_ ← L(t)*. (cell division)

## Acknowledgements

We thank Prof. Erin O'Shea for the Δ*kaiBC* plasmid, and Katie Abley and Henrik Jönsson for critical reading of the manuscript and useful suggestions. This research was made possible by the award of a European Research Council under the European Union’s Seventh Framework Programme (FP/2007-2013)/ERC Grant Agreement 338060. The work in the Locke laboratory is further supported by a fellowship from the Gatsby Foundation (GAT3272/GLC) and a Fellowship from the Human Frontier Science Program (CDA00068/2012). BMCM is supported by the UK Biotechnological and Biological Sciences Research Council (BBSRC) Synthetic Biology Research Centre “OpenPlant” award (BB/ L014130/1). PT gratefully acknowledges support by The Royal Commission for the Exhibition of 1851.

## Supporting Information

### 1 Segmentation and cell length

All images were segmented and tracked using a modified version of Schnitzcells (1). The segmentation algorithm is performed on a combined fluorescent-phase contrast image. The fluorescent image is acquired in the range of red wavelengths, and so it collects the cells' auto-fluorescence. Since *S. elongatus* cells are rod shaped, cell size is proportional to cell length. Cell length is defined as the length of the semi-major axis of the segmented cell shapes.

### 2 Elongation rates

Time-series of cell length were extracted and smoothed with either a locally weighted regression (*lowess*) method (for constant light data) or a moving average window 3 data points wide (LD cycles). We defined elongation rate as

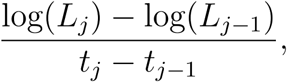

where *L_j_* is the smoothed cell length at time *t_j_*.

### 3 Defining subjective time of birth and division under constant light

Cells were entrained by 12:12 LD cycles (step cycles) before acquisition. We consider subjective dawn to occur immediately after the lights are switched on following the last 12 h long dark pulse. Times of day relative to a 24-hour day are obtained by rescaling total experimental time *t_exp_* by *t_exp_* modulo 24 hours. Accordingly, subsequent subjective dawn times are considered to occur at multiples of 24 h later. We define subjective dusk times to occur 12 h after subjective 20dawn.

### 4 Subpopulation clustering

For each tracked single cell that completed a cell cycle during image acquisition, we extracted distributions of time of birth, time of division, length at birth, length at division, added length and cell cycle time. Joint distributions of time of birth and cell cycle time revealed two distinct peaks (Fig. 2C). We extracted the cells that lie in each subpopulation by clustering with a two-component Gaussian mixture model using a likelihood-ratio criterion.

### 5 Analysis of division times using lineage-weighting

To analyse the frequency of cell divisions throughout the day, we used a lineage-weighted kernel density estimator in Fig. 6B,D. To this end, we extracted all lineages from the data and weighted each data point by a factor 1/2*^D^*depending on the number of cell divisions *D* in that lineage. The method takes into account that fast-growing cells are overrepresented compared to slower growing ones (2, 3). We found the difference to the unweighted data is significant only in the LD conditions, because the two subpopulations have vastly different interdivision times.

### 6 Estimating cell size control in the clock-deletion strain

We parameterized the dynamic model of the size control of the clock-deletion strain by first estimating the slope *a* through a linear regression. We then estimate the residuals of the regression via Δ_0_ = *L* − (1 + *a*) *L*_0_, and fit these data by a Gamma distribution *p*(Δ_0_). The size control hazard *S*(Δ_0_) is obtained as the ratio between this distribution and its survival function, giving the relation

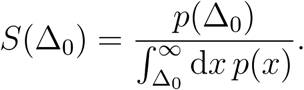

For a Gamma distribution with shape and scale parameters α and β, the above evaluates to

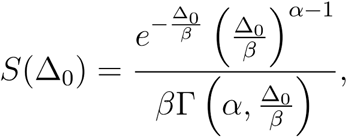

where Γ denotes the upper incomplete gamma function. This procedure was used to fit the size control in Fig. 3-6 from the corresponding experiments of clock-deletion cells (see Supp. Fig. 7 for parameters).

### 7 Inference of the circadian coupling function from single-cell data

To determine the coupling function, we need to estimate its likelihood. The probability for a single cell born at time *t*_0_ to divide after a cell cycle time τ is a product of the probability of cell division 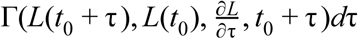 and the probability that the given cell has not divided before that time. The result is

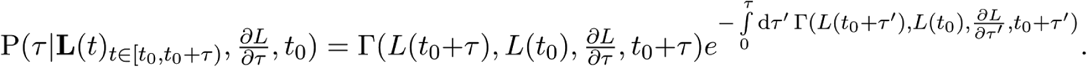

This expression depends not only on the whole single-cell trajectory of length but also on its derivative with respective to cell age τ^′^.

Because derivatives are difficult to estimate from single-cell data, we focus on the likelihood for a cell to divide at a specific length, which can be obtained by a change of variable. Using Eq. (1) in the equation above, we obtain the probability of a cell to divide at a length *L*,

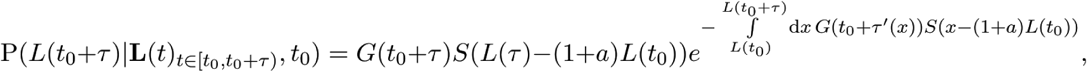

a quantity that is independent of the length derivatives but depends on cell age τ ′(*L*). Since cell length strictly increases in almost all of the observed single-cell traces, we estimated this quantity by interpolating cell age against cell length measurements for each cell using a cubic B-spline and evaluated the resulting integral in the above equation numerically.

The likelihood of the coupling function given *N* single-cell observations is then given by

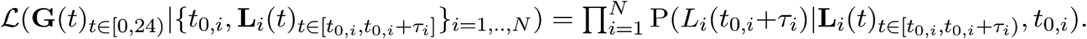

We parameterized the coupling function *G*(*t*) = *exp*(*B*(*t*)), a positive function with arguments taken modulo 24 hours, by a cubic B-spline *B*(*t*) with 12 knots (0,2,…,22 h) and periodic boundary conditions. The spline was evaluated at the knots and we sampled the posterior distributions using an adaptive Gibbs-sampler implemented in the Julia library *Mamba* (4). The result of this inference is shown in Fig. 3C.

## Supplementary figures

**Supplementary Figure 1:**
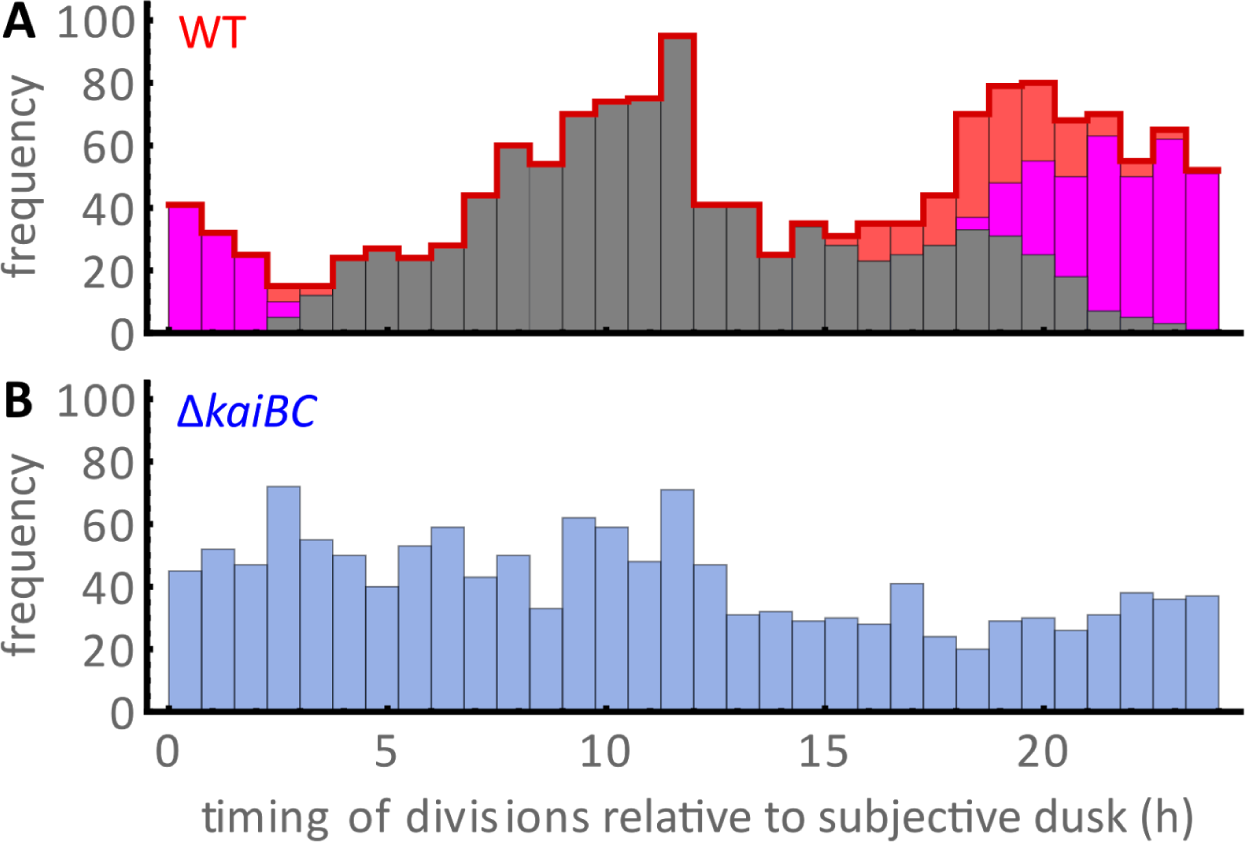
Distribution of division times in constant light. **(A)** Division times in WT are non-uniform, displaying a bimodal distribution (red). Clustering of subpopulations shows that fast cells (magenta histogram) divide mostly before dusk at the end of the subjective day while slow cells (grey histogram) divide around subjective dawn. **(B)** In the clock mutant (bottom) the distribution is more uniform. Sample sizes as in Fig. 2.

**Supplementary Figure 2:**
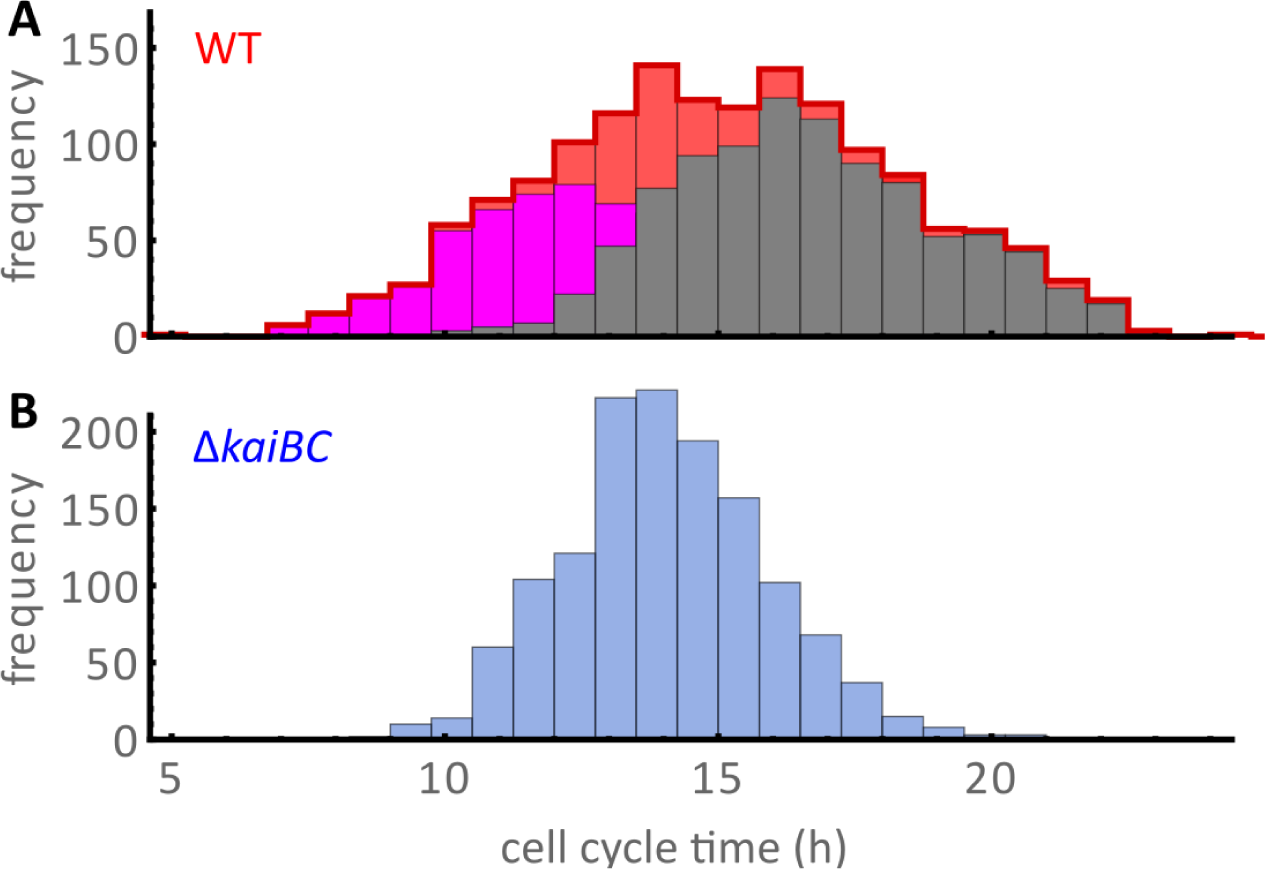
Distribution of cell cycle times in constant light. **(A)** Histograms show overlapping, but distinct, subpopulations with shorter (magenta) and longer cell cycle times (grey), respectively. The overall distribution is only weakly bimodal (red). **(B)** In clock-deletion cells, the histogram of cell cycle times is unimodal and narrower than in the wild type. Sample sizes as in Fig. 2.

**Supplementary Figure 3:**
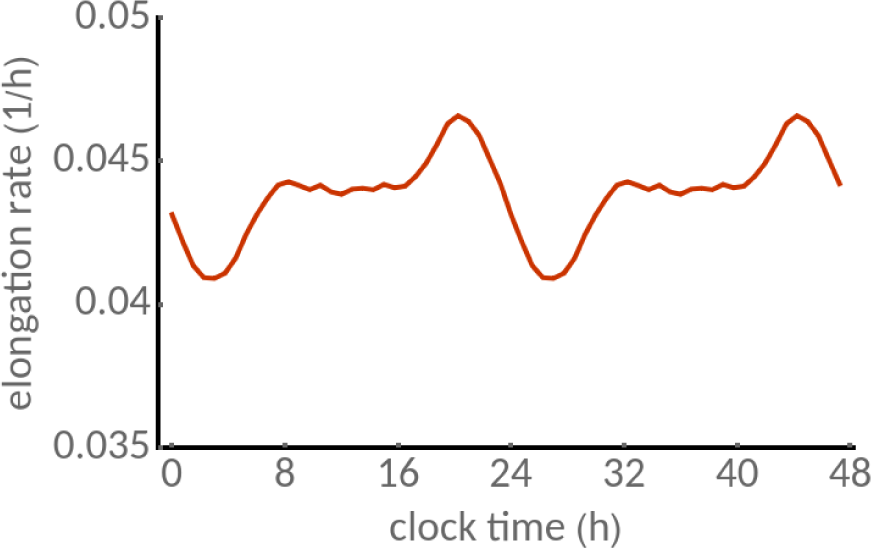
Mean elongation rate in constant light experiments. WT elongation rates oscillate throughout the day with a circadian period. Elongation rate peaks before subjective dusk and has a trough in early subjective night. Data replicated for two cycles to highlight periodicity.

**Supplementary Figure 4:**
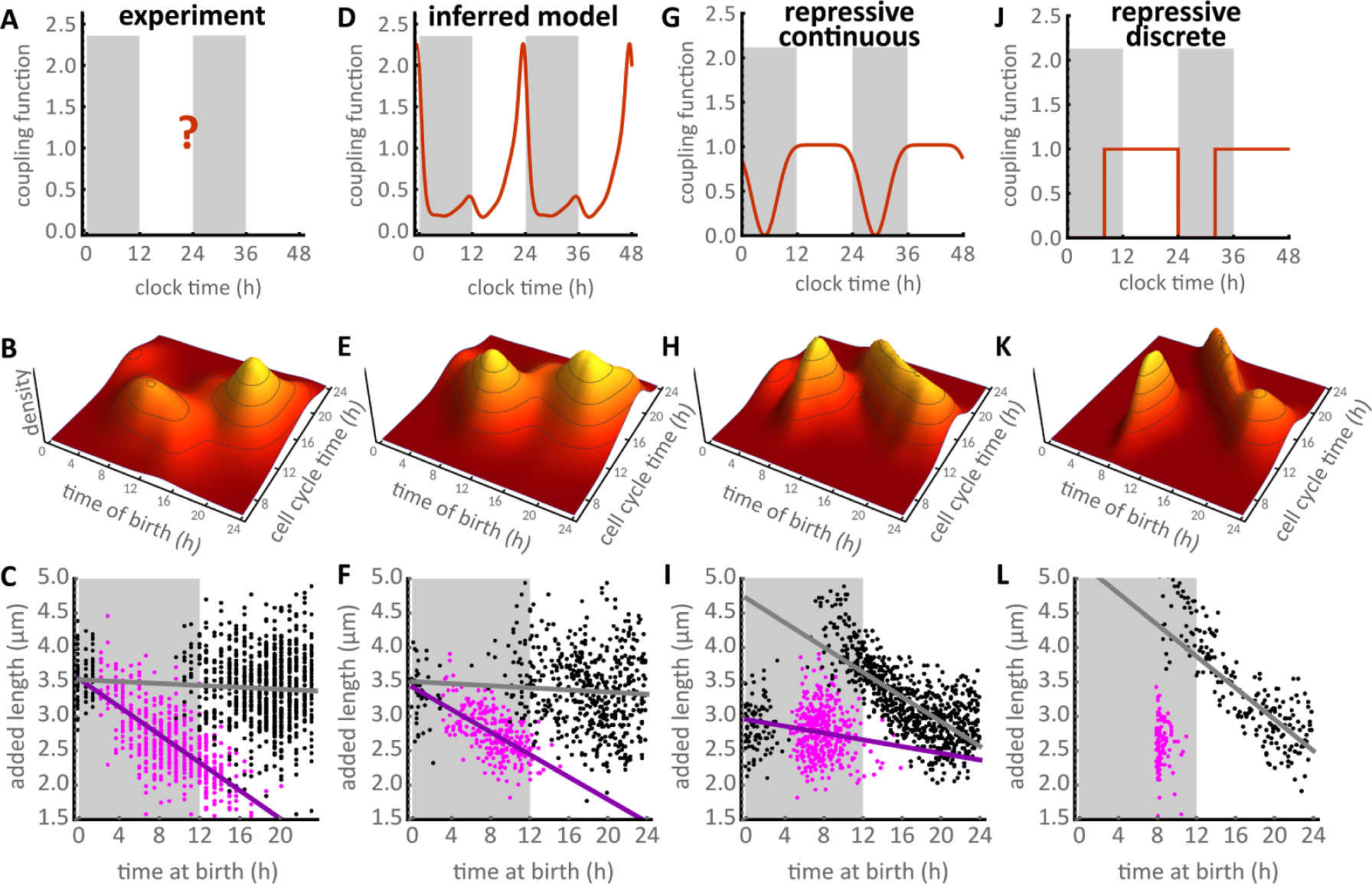
Cell size control identifies the coupling function. **(A-C)** Real coupling function is unknown. Experimental cell cycle times display slow and fast subpopulations of cells born at different times of the 24-h day. Added length decreases throughout the day in the fast subpopulation (magenta dots) but not in the slow one (black dots). **(D-F)** Coupling function inferred in this study, which promotes cell divisions towards the end of subjective day. This coupling correctly predicts the dependence of added length on time of birth. **(G-J)** Two repressive couplings (on-off gates) proposed in the literature inhibit cell divisions in the beginning of subjective night. These couplings also generate two or three subpopulations but display decreasing added lengths in the slow subpopulations and a lower slope (violet lines) in the fast ones, which is not supported by our experiments (C). In the lower panels, magenta and black dots are single cell data points, and violet and grey lines are the regression lines for the fast and slow subpopulations, respectively.

**Supplementary Figure 5:**
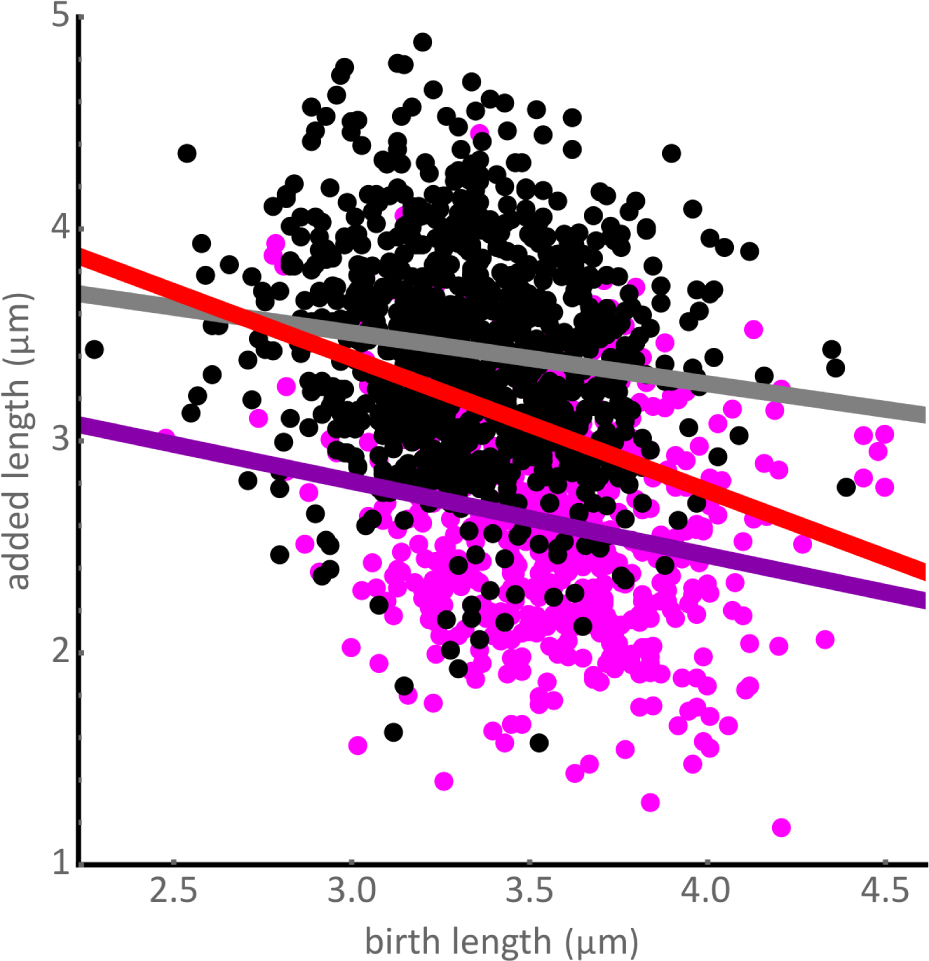
Sizer-like principle in WT cells in constant light is a consequence of ignoring two subpopulations. The two subpopulations of cells (magenta and black dots) add different lengths. The slopes characterising the dependence of added length on birth length of each subpopulation (violet and grey lines with slopes −0.36 and −0.25, respectively) are less steep than the global slope (red line, slope −0.62).

**Supplementary Figure 6:**
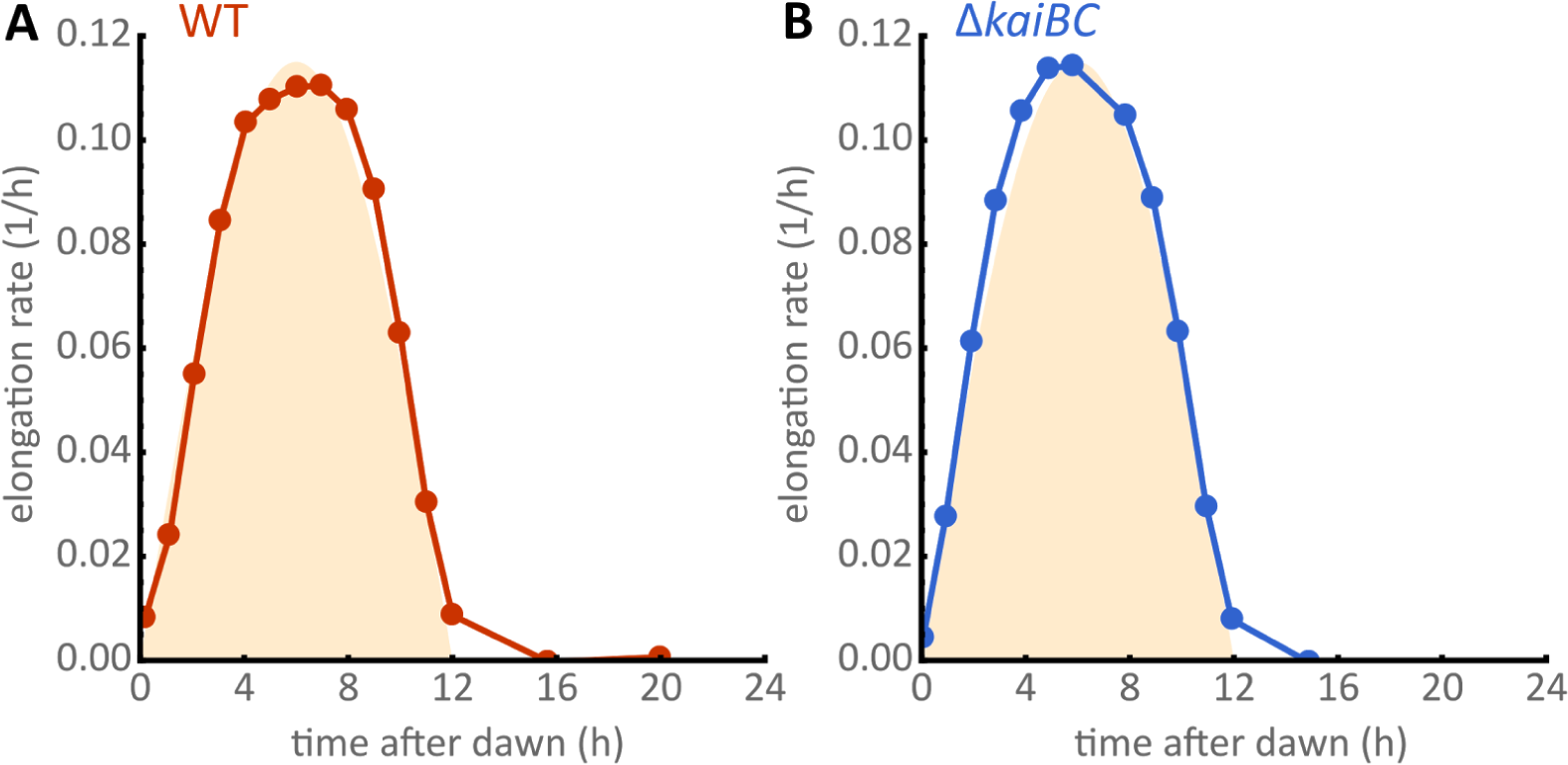
Exponential elongation rates in 12:12 LD cycles. **(A)** The mean elongation rate of WT cells (red dots) in a 24 h period closely follows the imposed light profiles (yellow shade). **(B)** The mean elongation rate of clock deletion cells (blue dots) is nearly identical to the WT under these conditions and is also determined by the light profile.

**Supplementary Figure 7:**
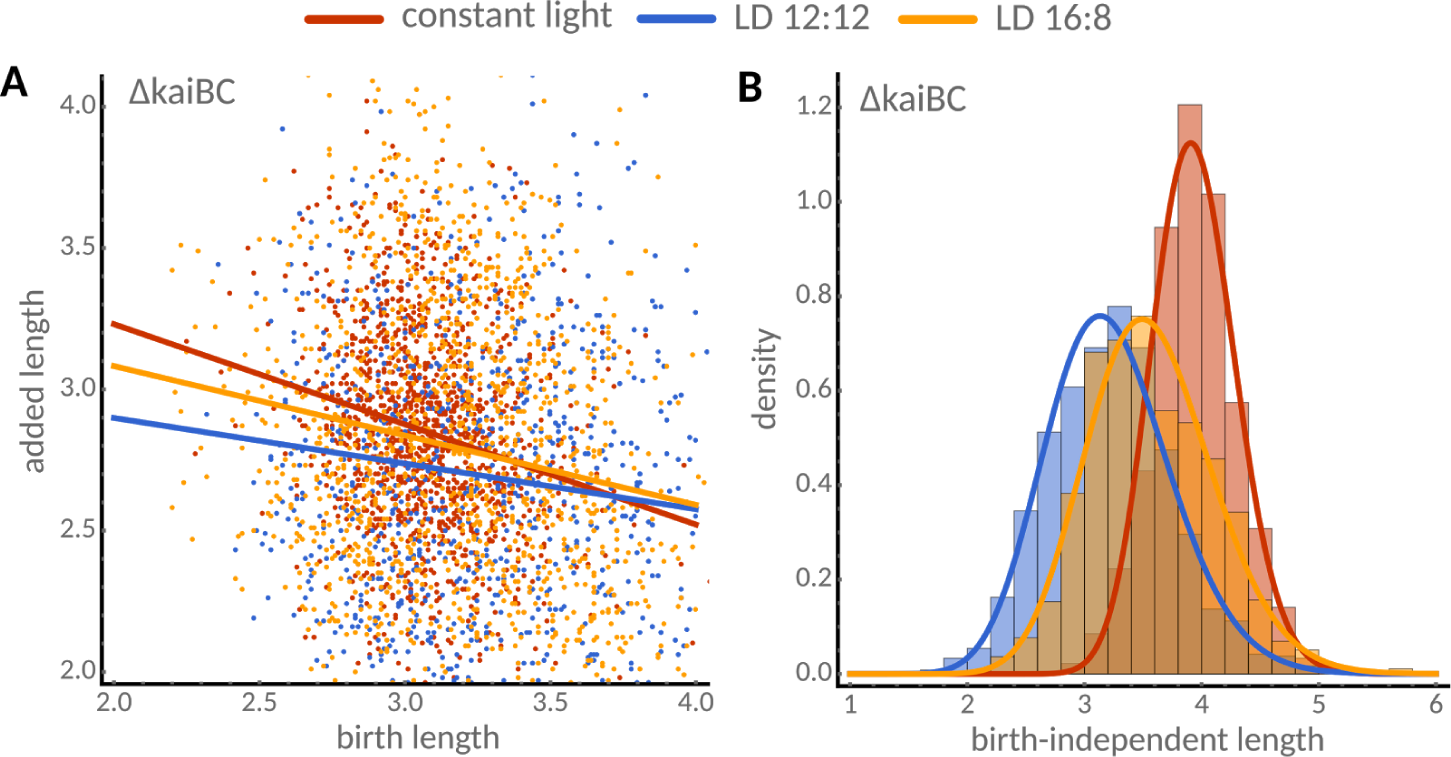
Characterisation of cell size control in clock-deletion cells. **(A)** Dependence of added length on birth length in constant light (red, slope −0.35), 12:12 (blue, slope −0.16) and 16:8 light-dark cycles (yellow, slope −0.25). **(B)** Distributions of birth-length independent part of added length, Δ0 = *L* − (1 + *a*) *L*0 (birth length *L*_0_, division length *L*, slope *a*, are well described by gamma distributions in all conditions. Fitted location and scale parameters are α = 122.5 and β = 0.03 (constant light), α = 36.6 and β = 0.09 (12:12 LD), and α = 44.4 and β = 0.08 (16:8 LD).

## References

1. Wood E, Nurse P (2015) Sizing up to divide: mitotic cell-size control in fission yeast. Annu Rev Cell Dev Biol 31:11–29.

2. Amodeo AA, Skotheim JM (2016) Cell-size control. Cold Spring Harb Perspect Biol 8(4):a019083.

3. Chien A-C, Hill NS, Levin PA (2012) Cell size control in bacteria. Curr Biol 22(9):R340–9.

4. Gonzalez N, Vanhaeren H, Inzé D (2012) Leaf size control: complex coordination of cell division and expansion. Trends Plant Sci 17(6):332–340.

5. Cooper S (2004) Control and maintenance of mammalian cell size. BMC Cell Biol 5(1):35.

6. Orskov J (1922) Method for the isolation of bacteria in pure culture from single cells and procedure for the direct tracing of bacterial growth on a solid medium https://doi.org/10.1101/175406. J Bacteriol 7(6):537–549.

7. Kelly CD, Rahn O (1932) The growth rate of individual bacterial cells. J Bacteriol 23(2):147–153.

8. Schaechter M, Maaloe O, Kjeldgaard NO (1958) Dependency on medium and temperature of cell size and chemical composition during balanced grown of Salmonella typhimurium. J Gen Microbiol 19(3):592–606.

9. Turner JJ, Ewald JC, Skotheim JM (2012) Cell size control in yeast. Curr Biol 22(9):R350–9.

10. Osella M, Nugent E, Lagomarsino MC (2014) Concerted control of Escherichia coli cell division. Proc Natl Acad Sci U S A 111(9):3431–3435.

11. Wang P, Hayden S, Masui Y (2000) Transition of the blastomere cell cycle from cell size-independent to size-dependent control at the midblastula stage in Xenopus laevis. J Exp Zool 287(2):128–144.

12. Sveiczer A, Novak B, Mitchison JM (1996) The size control of fission yeast revisited. J Cell Sci 109 (Pt 12):2947–2957.

13. Nobs J-B, Maerkl SJ (2014) Long-term single cell analysis of S. pombe on a microfluidic microchemostat array. PLoS One 9(4):e93466.

14. Iyer-Biswas S, et al., (2014) Scaling laws governing stochastic growth and division of single bacterial cells. Proc Natl Acad Sci U S A 111(45):15912–15917.

15. Sauls JT, Li D, Jun S (2016) Adder and a coarse-grained approach to cell size homeostasis in bacteria. Curr Opin Cell Biol 38:38–44.

16. Campos M, et al., (2014) A constant size extension drives bacterial cell size homeostasis. Cell 159(6):1433–1446.

17. Amir A (2014) Cell Size Regulation in Bacteria. Phys Rev Lett 112(20).

18. Taheri-Araghi S, et al., (2015) Cell-size control and homeostasis in bacteria. Curr Biol 25(3):385–391.

19. Soifer I, Robert L, Amir A (2016) Single-cell analysis of growth in budding yeast and bacteria reveals a common size regulation strategy. Curr Biol 26(3):356–361.

20. Deforet M, van Ditmarsch D, Xavier JB (2015) Cell-size homeostasis and the incremental rule in a bacterial pathogen. Biophys J 109(3):521–528.

21. Yu FB, et al. (2017) Long-term microfluidic tracking of coccoid cyanobacterial cells reveals robust control of division timing. BMC Biol 15(1):11.

22. Kuwano K, et al. (2014) Growth and cell cycle of Ulva compressa (Ulvophyceae) under LED illumination. J Phycol 50(4):744–752.

23. Bišová K, Zachleder V (2014) Cell-cycle regulation in green algae dividing by multiple fission. J Exp Bot 65(10):2585–2602.

24. Nishihama R, Kohchi T (2013) Evolutionary insights into photoregulation of the cell cycle in the green lineage. Curr Opin Plant Biol 16(5):630–637.

25. Nozue K, et al., (2007) Rhythmic growth explained by coincidence between internal and external cues. Nature 448(7151):358–361.

26. Doherty CJ, Kay SA (2010) Circadian control of global gene expression patterns. Annu Rev Genet 44:419–444.

27. Sweeney BM, Woodland Hastings J (1958) Rhythmic cell division in populations of Gonyaulax polyedra. J Protozool 5(3):217–224.

28. Mori T, Binder B, Johnson CH (1996) Circadian gating of cell division in cyanobacteria growing with average doubling times of less than 24 hours. Proc Natl Acad Sci U S A 93(19):10183–10188.

29. Bieler J, et al., (2014) Robust synchronization of coupled circadian and cell cycle oscillators in single mammalian cells. Mol Syst Biol 10:739.

30. Feillet C, et al., (2014) Phase locking and multiple oscillating attractors for the coupled mammalian clock and cell cycle. Proc Natl Acad Sci U S A 111(27):9828–9833.

31. Matsuo T, et al., (2003) Control mechanism of the circadian clock for timing of cell division in vivo. Science 302(5643):255–259.

32. Kuan D, Duff S, Posarac D, Bi X (2015) Growth optimization of Synechococcus elongatus PCC7942 in lab flasks and a 2-D photobioreactor. Can J Chem Eng 93(4):640–647.

33. Chabot JR, Pedraza JM, Luitel P, van Oudenaarden A (2007) Stochastic gene expression out-of-steady-state in the cyanobacterial circadian clock. Nature 450(7173):1249–1252.

34. Dong G, et al., (2010) Elevated ATPase activity of KaiC applies a circadian checkpoint on cell division in Synechococcus elongatus. Cell 140(4):529–539.

35. Teng S-W, Mukherji S, Moffitt JR, de Buyl S, O’Shea EK (2013) Robust circadian oscillations in growing cyanobacteria require transcriptional feedback. Science 340(6133):737–740.

36. Rust MJ, Markson JS, Lane WS, Fisher DS, O’Shea EK (2007) Ordered phosphorylation governs oscillation of a three-protein circadian clock. Science 318(5851):809–812.

37. Cohen SE, Golden SS (2015) Circadian rhythms in cyanobacteria. Microbiol Mol Biol Rev 79(4):373–385.

38. Ito H, et al., (2007) Autonomous synchronization of the circadian KaiC phosphorylation rhythm. Nat Struct Mol Biol 14(11):1084–1088.

39. Markson JS, Piechura JR, Puszynska AM, O’Shea EK (2013) Circadian control of global gene expression by the cyanobacterial master regulator RpaA. Cell 155(6):1396–1408.

40. Diamond S, Jun D, Rubin BE, Golden SS (2015) The circadian oscillator in Synechococcus elongatus controls metabolite partitioning during diurnal growth. Proc Natl Acad Sci U S A 112(15):E1916–25.

41. Pattanayak GK, Phong C, Rust MJ (2014) Rhythms in energy storage control the ability of the cyanobacterial circadian clock to reset. Curr Biol 24(16):1934–1938.

42. Yang Q, Pando BF, Dong G, Golden SS, van Oudenaarden A (2010) Circadian gating of the cell cycle revealed in single cyanobacterial cells. Science 327(5972):1522–1526.

43. Lambert G, Chew J, Rust MJ (2016) Costs of Clock-Environment Misalignment in Individual Cyanobacterial Cells. Biophys J 111(4):883–891.

44. Tanouchi Y, et al., (2015) A noisy linear map underlies oscillations in cell size and gene expression in bacteria. Nature 523(7560):357–360.

45. Priestman M, Thomas P, Robertson BD, Shahrezaei V (2017) Mycobacteria modify their cell size control under sub-optimal carbon sources. Front Cell Dev Biol 5. doi:10.3389/fcell.2017.00064.

46. Modi S, Vargas-Garcia CA, Ghusinga KR, Singh A (2017) Analysis of Noise Mechanisms in Cell-Size Control. Biophys J 112(11):2408–2418.

47. Martins BM, Das AK, Antunes L, Locke JC (2016) Frequency doubling in the cyanobacterial circadian clock. Mol Syst Biol 12(12):896.

48. Leypunskiy E, et al. (2017) The cyanobacterial circadian clock follows midday in vivo and in vitro. Elife 6. doi:10.7554/eLife.23539.

49. Mori T, Johnson CH (2001) Independence of circadian timing from cell division in cyanobacteria. J Bacteriol 183(8):2439–2444.

50. Young JW, et al., (2011) Measuring single-cell gene expression dynamics in bacteria using fluorescence time-lapse microscopy. Nat Protoc 7(1):80–88.

51. Lewis PA, Shedler GS (1979) Simulation of Nonhomogeneous Poisson Processes by Thinning. Nav Res Logist 26:403–413.

52. Voliotis M, Thomas P, Grima R, Bowsher CG (2016) Stochastic Simulation of Biomolecular Networks in Dynamic Environments. PLoS Comput Biol 12(6):e1004923.

## References

1 Young JW, et al., (2011) Measuring single-cell gene expression dynamics in bacteria using fluorescence time-lapse microscopy. Nat Protoc 7(1):80–88.

2 Nozoe T, Kussell E, Wakamoto Y (2017) Inferring fitness landscapes and selection on phenotypic states from single-cell genealogical data. PLoS Genet 13(3):e1006653

3. Priestman M, Thomas P, Robertson BD, Shahrezaei V (2017) Mycobacteria modify their cell size control under sub-optimal carbon sources. Front Cell Dev Biol 5. doi:10.3389/fcell.2017.00064.

4. Smith BJ et al Mamba: Markov chain Monte Carlo for Bayesian analysis in Julia Available at: https://github.com/brain-j-smith/Mamba.jl.

